# Bridging the gap between *in silico* and *in vivo:* modeling opioid disposition in a kidney proximal tubule microphysiological system

**DOI:** 10.1101/2020.12.04.412239

**Authors:** Tomoki Imaoka, Weize Huang, Sara Shum, Dale W. Hailey, Shih-Yu Chang, Alenka Chapron, Catherine K. Yeung, Jonathan Himmelfarb, Nina Isoherranen, Edward J. Kelly

## Abstract

**Background:** Opioid overdose, dependence, and addiction are a major public health crisis. Patients with chronic kidney disease (CKD) are at high risk of opioid overdose, therefore novel methods that provide accurate prediction of kidney clearance (CL_r_) and systemic disposition of opioids in CKD patients can facilitate the optimization of therapeutic regimens.

**Methods:** We conducted prediction of kidney clearance and systemic disposition of morphine and its active metabolite morphine-6-glucuronide (M6G) in CKD patients using a vascularized human proximal tubule microphysiological system (VPT-MPS) coupled with a parent-metabolite full body physiologically-based pharmacokinetic (PBPK) model.

**Results:** The VPT-MPS, populated with a human umbilical vein endothelial cell (HUVEC) channel and an adjacent human primary proximal tubular epithelial cells (PTEC) channel, successfully demonstrated secretory transport of morphine and M6G from the HUVEC channel into the PTEC channel in a time-dependent manner; transporter inhibitors decreased translocation by 74.3% and 63.6%, respectively. The *in vitro* data generated by VPT-MPS were incorporated into a mechanistic kidney model and parent-metabolite full body PBPK model to predict CL_r_ and systemic disposition of morphine and M6G. The model successfully predicted CL_r_ within 1.5-fold, and the plasma concentration-time profiles of morphine and M6G in both healthy subjects and CKD patients, with absolute average fold error values <1.5.

**Conclusions:** A microphysiological system together with mathematical modeling successfully predicted kidney clearance and systemic disposition of opioids in CKD patients and healthy subjects.

## Introduction

The kidney plays a pivotal role in the elimination of xenobiotics and their metabolites from the body, affecting drug efficacy and safety profiles. Kidney excretion is achieved by the coordinated actions of glomerular filtration, transporter-mediated active secretion, and passive reabsorption. Accordingly, in order to evaluate kidney drug excretion *in vitro*, a model must incorporate both transporter-mediated active secretion and passive permeation processes across the tubular epithelium. Several methodologies have been proposed to predict active kidney secretion of drugs including kidney slices (1), *in vitro*-*in vivo* extrapolation (IVIVE) approaches using transfected cells (2), and PTEC monolayers in 2D Transwell™(3). However, these methodologies have limitations that may restrict their utility. For example, kidney slices fail to evaluate net secretion of drugs across proximal tubule lumen due to the occlusion of cut ends of proximal tubules (4). IVIVE approaches usually assume that the tubular secretion is determined by only OAT-mediated transport and do not consider other transporters such as OCT2, OATP4C1 or unknown transporter(s) that may also contribute to net kidney secretion (2). Although PTEC monolayers in 2D Transwell™ may help evaluate kidney secretion of drugs by evaluation of directional transport, our previous investigations failed to observe active transport of probe substrates of kidney transporters (5) presumably because the lack of fluid shear stress in the Transwell™ system results in the loss of transporter expression (6).

In light of these observations, there is increasing interest in organ-on-chips technology as an improved system due to the incorporation of physiomimetic flow and shear stress (7). Several research teams have generated ‘kidney-on-a chip’ microphysiological systems (MPS) populated with PTECs that successfully recapitulate key component structures and functions of the human proximal tubule (8). Although PTECs subjected to physiological tubular flow exhibit enhanced differentiation as revealed by increased primary cilium formation, alkaline phosphatase activity, albumin transport, and glucose reabsorption, compelling data demonstrating secretory transport of organic solutes is lacking. Recently our group has, for the first time, observed secretory clearance of organic solutes in a proximal tubule MPS (5). This prompted further development of a vascularized human proximal tubule microphysiological system (VPT-MPS), where PTEC and HUVEC tubules are created in adjacent channels to evaluate secretory function of the kidney. This platform is expected to be a useful tool to define kidney disposition characteristics in human as it enables quantitation of transporter expression and evaluation of tubular secretion of xenobiotics and endogenous compounds (e.g., *para*-aminohippuric acid (9)).

In addition to the technical difficulties involved in the *in vitro* evaluation of kidney secretory clearance, prediction of kidney clearance must also quantitatively account for complex kidney physiology and dynamic kidney drug handling. As such, we have developed and verified a dynamic physiologically-based mechanistic kidney model that incorporates unbound filtration, active secretion, and tubular flow rate- and filtrate pH-dependent passive reabsorption (10). Our hypothesis is that incorporating the *in vitro* data generated from the VPT-MPS into the physiologically-based mechanistic kidney model will allow prediction of CL_r_ of morphine and M6G in humans. Further, using the mechanistic kidney model-integrated full body parent-metabolite PBPK model (11), plasma concentration-time profile of morphine and M6G can be predicted in healthy subjects. Finally, due to the increased risk of opioid overdose in people with CKD (12-14), the developed model can be extrapolated (15) to simulate systemic disposition of morphine and M6G in varying stages of CKD. This is the first study to demonstrate the combination of a VPT-MPS and mechanistic PBPK modeling for the quantitative prediction of kidney handling and plasma disposition of drugs in healthy subjects and people with CKD.

## Methods

### Chemicals and reagents

Morphine (1.0 mg/mL, in methanol solution), morphine-d6 (1.0 mg/mL, in methanol solution), morphine-6-beta-D-glucuronide (1.0 mg/mL, in methanol solution), bovine serum albumin, hydrocortisone, Triton X-100, probenecid and tetraethylammonium were purchased from Sigma-Aldrich (St. Louis, MO). ^3^H-PAH (40 Ci/mmol) was obtained from American Radiolabeled Chemicals (St. Louis, MO). Formaldehyde (16%, methanol free) was purchased from Polysciences (Warrington, PA). Phosphate-buffered saline with (PBS), 50:50 Dulbecco’s modified eagle medium with Ham’s F-12 (DMEM/F12), penicillin-streptomycin-amphotericin B, insulin-transferrin-selenium A solution (ITS-A), fetal bovine serum (FBS), 0.05% Trypsin EDTA, ProLong Gold Antifade reagent were purchased from Thermo-Fisher (Waltham, MA). Alexa Fluor 594 conjugated anti-mouse IgG, Alexa Fluor 488 conjugated anti-rabbit IgG, and anti-OAT3 antibody were obtained from Abcam (Cambridge, MA). Anti-OAT1 antibody was obtained from Fischer Scientific (Pittsburgh, PA). Anti-OCT2 antibody was obtained from RD systems (Minneapolis, MN). Dual-channel MPS platforms were supplied by Nortis (Woodinville, WA). Rat tail collagen I was purchased from Ibidi (Martinsried, Germany). Collagen IV was obtained from Corning (Corning, NY). Human umbilical venous endothelial cells (HUVECs) and EGM-2 bullet kits were obtained from Lonza (Basel, Switzerland).

### Isolation of PTECs

Human PTECs were isolated as previously described (5), in accordance with a protocol approved by the University of Washington Human Subjects Institutional Review Board (# STUDY00001297). Detailed kidney tissue donor information is available in Supplementary Table S1. Following isolation, cells were expanded in tissue culture flasks to passages 1-2 before use in all experiments.

### Establishment of vascularized proximal tubule MPS (VPT-MPS)

The inner chamber of dual-channel MPS devices (Duplo™, Figure 1) (https://www.nortisbio.com/) was dried, filled with 6 mg/mL of rat tail collagen I, and left overnight to allow the collagen I matrix to solidify. The mandrels of the MPS platforms were then removed, leaving two hollow channels through the collagen I matrix. The PTEC channel was coated with a 5 µg/mL mouse collagen IV to facilitate human PTECs adhesion. Cultures of primary kidney epithelial cells were subjected to trypsin digestion to obtain single cell suspensions. The cells were counted and resuspended with PTECs culture media at a concentration of 20 ×10^6^ cells/mL and 2-2.5 µL were injected into the collagen IV-coated channel. Cells were allowed to attach for 3 hours before starting perfusion at a rate of 1.0 µL/min. A 1.0 µL/min perfusion of DMEM/F12 media containing ITS-A, 50 nM hydrocortisone, penicillin-streptomycin and amphotericin B was then initiated. Following 6 days under these culture conditions, the epithelial tubule was established and perfusion to the channel was stopped for a brief period. The parallel vacant channel was equilibrated with EGM-2 media containing 2 % FBS for 2 hours at 1.0 µL/min. Then, HUVECs (passages 5 to 10) were seeded into the equilibrated channel and allowed to attach for 30 minutes to establish an endothelial vessel. Flow to both channels was resumed at 1.0 µL/min with both channels receiving their respective culture medium. The day after seeding HUVEC, VPT-MPS were used for transport experiments. Prior to the beginning of experiments, a visual inspection under light microscope was performed to confirm complete (100%) coverage of both HUVEC vessel and PTEC tubule, respectively. Only VPT-MPS that exhibited even, non-perturbed media flow characteristics (i.e., nearly identical volume of output media from HUVEC vessel and PTEC tubule) were selected for use in functional experiments.

**Figure. 1.**
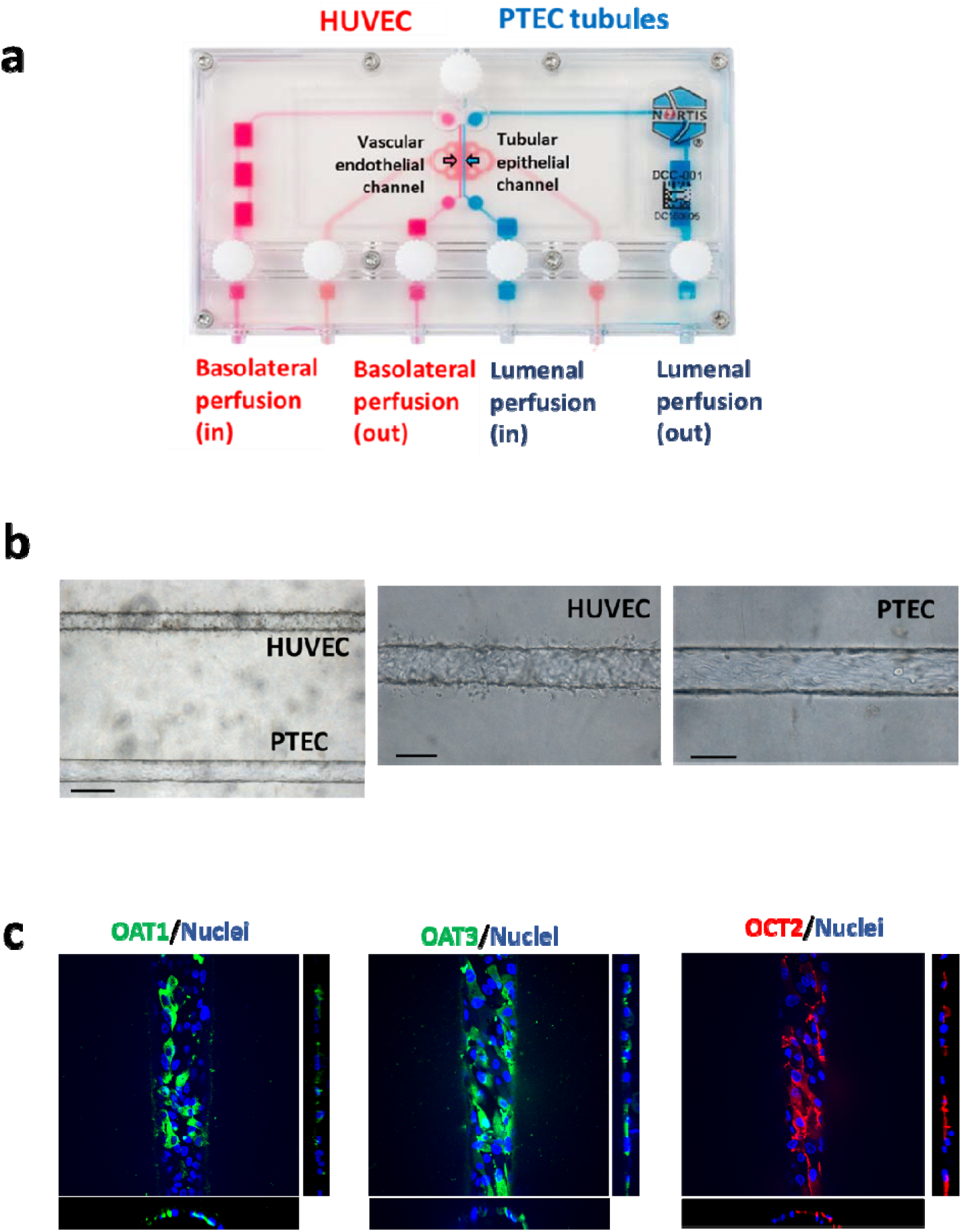
Establishment of vascularized proximal tubule MPS (VPT-MPS) (a) Nortis dual channel MPS devices (Duplo™) used for the establishment of vascularized proximal tubule MPS (VPT-MPS). Vascular media flow path is shown in pink and tubular media flow is shown in blue. (b) Bright field images of the bioengineered co-culture in the collagen I matrix by 10x (left) and 20x magnification (middle and right). PTECs were seeded and allowed to grow for 6 days, followed by seeding of HUVECs for 24 hours prior to transport experiment. (c) Determination of transporter expression in PTECs via immunocytochemistry. Top down maximum intensity projection reveals expression of OAT1, OAT3 and OCT2 in the cells in the channel; side and lower panels show XZ and YZ cross sections, where X is along the length of the channel and Y is across the curvature. Cells expressed OAT1 (left), OAT3 (middle), and OCT2 (right). Nuclei are shown in blue. Negative control (treated with only secondary antibody) showed minimal fluorescence (data not shown). Average diameter of the tubule is 120 µm (200x magnification). Representative images of cells isolated from a single donor.

### Immunocytochemistry

Vascularized proximal tubule MPS were fixed with a 1 hr perfusion of 4% formaldehyde in PBS++ through both channels at 1 uL/min, followed by a wash with PBS++ (containing calcium and magnesium) at a rate of 5 uL/min for 30 minutes. Blocking of non-specific binding was achieved by exposing the cells to a solution of 0.1% Triton X-100 and 5% bovine serum albumin in PBS++ (PTB). Primary antibodies, diluted in PTB, were then introduced into the channels at the following concentrations: rabbit OAT1 1:20, rabbit OAT3 1:100, mouse OCT2 1:20, (incubation time: 16 h at 4° C). Following a wash with PBS++, secondary antibodies, diluted 1:1000 in PTB, were perfused (incubation time: 1-2 h at room temperature). The cells were again washed with PBS++ and then finally exposed to a perfusion of a mixture of 1:4 dilution of ProLong Gold Anti-fade with 2 droplets of NucBlue for staining cell nuclei. Negative controls consisted of perfused samples that were not incubated with primary antibodies. The vascularized proximal tubule MPS platforms were imaged with a Yokogawa W1 spinning disk confocal system on a Nikon Eclipse Ti inverted microscope with a CFI60 Apochromat Lambda S LWD 40x water immersion objective (N.A. 1.15) and Andor iLife EMCCD camera. Images were processed in ImageJ.

### Transport of morphine and M6G in the VPT-MPS

EGM-2 media containing morphine-d6 (1 µM) and M6G (1 µM) was perfused through the endothelial vessel, mimicking solute delivery via peritubular capillaries in the presence or absence of competitive inhibitors for transporters (mixture of probenecid (1mM) and tetraethylammonium (1mM)). The flow rate of flow through both channels was set at 1 µL/min. Effluents from the vessel and the tubule were collected every 2 hours, and drug concentrations in the samples were determined LC/MS/MS. Transport experiments were conducted across 3 different donors. Effluent measurements were extended out to 24 hours in order to reach steady-state levels in solute output. Clearances of morphine and M6G in VPT-MPS were calculated by following equation:

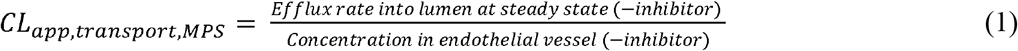

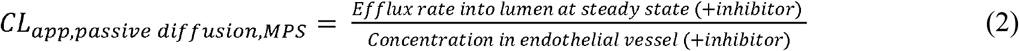

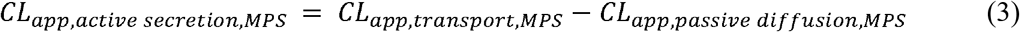

where CL_app, transport,MPS_, CL_app, passive diffusion,MPS_ and CL_app, active secretion,MPS_ represents net efflux clearance, permeability and active transporter-mediated secretion of morphine or M6G across PTECs tubules, respectively.

### Quantification of morphine and morphine-6-glucuronide using LC-MS/MS

Morphine and morphine-6-glucuronide concentrations in VTS-MPS and Transwell™ permeability samples were measured by an optimized HPLC-MS/MS method. The samples (50 µL) were thawed at room temperature on the day of analysis. For VPT-MPS samples, calibration standards were prepared in PTEC culture media (DMEM/F12 media containing added ITS-A, 50 nM hydrocortisone, penicillin, streptomycin and amphotericin B) with spiked morphine-d_6_ and morphine-6-glucuronide at concentrations ranging between 2 and 1,000 nM. For each sample or calibration standard, 100 µL of methanol containing 100 nM of morphine and morphine-6-glucuronide-d_3_ were added. For Transwell™ permeability samples, calibration standards were prepared in HBSS buffer with spiked morphine and morphine-6-glucuronide at concentrations ranging between 1 and 300 nM. For each sample or calibration standard, 100 µL of methanol containing 100 nM of morphine-d_6_ and morphine-6-glucuronide-d_3_ were added. The samples and standards were vortexed briefly and centrifuged at 3,000 g for 30 minutes at 4°C. Clear supernatant was removed and analyzed using HPLC-MS/MS using AB Sciex 6500 QTRAP system (Foster City, CA) connected in line with a Shimadzu UFLC XR DGU-20A5 (Kyoto, Japan) equipped with a Phenomenex Kinetex® EVO C18 (2.6 µm, 100 x 2.1 mm^2^) LC column and a guard cartridge (2 x 2.1 mm^2^, sub 2 µm) (Torrance, CA). Two microliters (2 µL) of supernatant was injected at each run at a flow rate of 0.35 mL/min. The elution gradient was initiated with 100% mobile phase A (10 mM ammonium formate) and 0% mobile phase B (50:50 v/v methanol and acetonitrile), increased to 10% B by 0.1 minute and continued at 10% B until 1 minute, then increased to 100% B in the next 3.5 minutes and continued at 100% B for 1 minute before returning to the initial condition in 0.1 minute and equilibrate for 3 minutes. Analytes were detected using electrospray ionization operated in positive ion mode. Three product ions for morphine (m/z 286 > 165, 286 > 181, 286 > 153), three product ions for morphine-d6 (m/z 292 > 165, 292 > 181, 292 > 153), one product ion for morphine-6-glucuronide (m/z 462 > 286) and one product ion for morphine 6-glucuronide-d3 (m/z 465 > 289) was monitored to confirm the identity of the analytes. Quantification of morphine and morphine-d6 were performed using the most abundant product ions m/z 286 > 165 and m/z 292 > 165, respectively. The lower limit of quantitation (LLOQ) of morphine, morphine-d6, and morphine-6-glucuronide are 2 nM.

### Prediction of kidney clearance of morphine and M6G using VPT-MPS data and mechanistic kidney model

The kidney clearance of morphine and M6G was simulated using the previously published and verified mechanistic kidney model (10) together with plasma unbound fraction (16, 17), blood-to-plasma ratio (18), and VPT-MPS-derived permeability and active secretion parameters. The permeability and active secretion parameters of morphine and M6G were derived as follows:

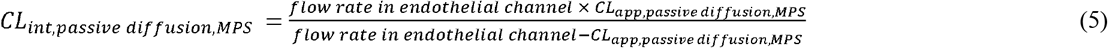

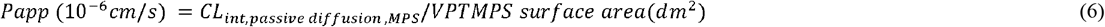

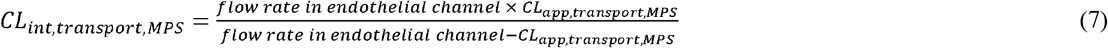

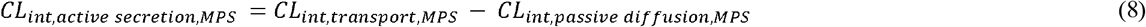

where CL_int, passive diffusion, MPS_ represents intrinsic passive diffusion clearance in the VPT-MPS; P_app_ represents the apparent transcellular permeability of morphine and M6G; CL_int, transport, MPS_ represents total intrinsic transport clearance in MPS; and CL_int, active secretion, MPS_ represents the intrinsic transporter-mediated active secretion clearance in the VPT-MPS.

To simulate the kidney clearance of morphine and M6G in human, the intrinsic active secretions obtained from VPT-MPS were further scaled up to the intrinsic secretion clearances in human (CL_secretion_) by multiplying with the 60 million proximal tubular cells per gram of kidney (19) and the kidney weight per human body (300 g) using the equation below.

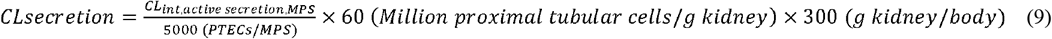

The final calculated permeability value, intrinsic active secretion, and the predicted kidney clearance of all three donors are summarized in Table 1.

**Table 1.** The summary of VPT-MPS derived intrinsic secretion clearance and permeability value for morphine and M6G, with predicted kidney clearance of morphine and M6G using mechanistic kidney model with VPT-MPS data

|  | Morphine |  |  | M6G |  |  |
| --- | --- | --- | --- | --- | --- | --- |
|  | donor1 | donor2 | donor3 | donor1 | donor2 | donor3 |
| $CL_{int,sec}$<br>($\mu L/hr/MPS$) | 14.1±11.9 | 1.33±0.57 | 20.87 | 15.6±10.5 | 1.29±1.09 | 22.9 |
| Permeability ($10^{-6}$<br>cm/s) | 27.8±22.3 | 13.9 | 42.7 | 25.9±28.5 | 9.74±11.1 | 41.8 |
| $CL_{r,predicted}$ (L/hr) | 9.69±5.18 | 4.78±0.61 | 8.27 | 11.6±3.14 | 7.23±1.61 | 9.49 |
| $CL_{r,observed}$ (L/hr) | 6.8-9.62<sup>(28, 33, 34)</sup> | | | 9.20-14.3<sup>(33, 34)</sup> | | |

### Development of morphine-M6G parent-metabolite full body PBPK model

To test whether the *in vitro* VPT-MPS morphine and M6G data could be used to simulate *in vivo* morphine and M6G kidney clearance and plasma concentration-time profiles in humans, a morphine-M6G parent-metabolite full body PBPK model was developed using MATLAB and Simulink platform (R2018a; MathWorks, Natick, MA). The structural model was established by merging our verified mechanistic kidney model (10) with the parent-metabolite PBPK model (20), as described previously (11). Specifically, this model incorporates unbound filtration, active secretion, and tubular flow rate and filtrate pH-dependent passive reabsorption to predict kidney drug clearance and thus allows the changes in mechanistic kidney clearance pathways to be reflected in systemic disposition. A schematic diagram of the model structure is shown in Figure 3. The drug models of morphine and M6G were developed independently and then linked together using a similar workflow and strategy as previously described by us for methamphetamine and amphetamine (11). The physicochemical properties of morphine and M6G were collected from literature (16-18, 21) and the permeabilities were determined using the VPT-MPS system. The distribution of morphine and M6G was governed by the tissue-to-plasma partition coefficients (K_p_) of 12 important tissues in the model. For morphine, the tissue-specific K_p_ values were first predicted using Rodger and Rowland’s method (22), resulting in a predicted V_ss_ of 285 L. As the observed V_ss_ from arterial morphine data is only 120 L (23) the tissue-specific K_p_ values for slow-perfusion tissues (adipose, muscle, and skin) were optimized (decreased to 0.5, 2.5, and 0.5, respectively), yielding a V_ss_ of 141 L. For M6G, it was assumed that M6G tissue distribution was limited mainly to blood and interstitial fluid given the observed V_ss_ values range from 19 to 29 L (24). Thus, a value of 0.3 was assigned to K_p_ values of all tissues to account for the space of interstitial fluid (extracellular water) of each tissue yielding a V_ss_ value of 21 L.

The systemic clearance of morphine and M6G included hepatic metabolism and kidney excretion in the model. The kidney clearance of morphine and M6G was simulated using the previously published and verified mechanistic kidney model (10) together with plasma unbound fraction (16, 17), blood-to-plasma ratio (18), and VPT-MPS-derived permeability and active secretion parameters. Hepatic clearance of morphine was obtained by subtracting the morphine kidney clearance from the observed morphine systemic clearance after intravenous administration (25). The intrinsic metabolic clearance of morphine was back-calculated based on its plasma unbound fraction, blood-to-plasma ratio, and the well-stirred hepatic clearance model (26). The M6G formation from morphine was modeled to occur within the liver compartment, and the fraction of metabolism (f_m_) from morphine to M6G was estimated as 10% based on literature (27). The hepatic clearance of M6G was assumed to be negligible. All the detailed physicochemical and pharmacokinetic values of morphine and M6G used in the model are summarized in Table 2.

**Table 2.** The summary of morphine-M6G parent-metabolite full body PBPK model parameters.

| Parameter | Morphine | M6G |
| --- | --- | --- |
| **Physicochemical** |  |  |
| Molecular weight (g/mol) | 285.34<sup>a</sup> | 461.46<sup>b</sup> |
| pK<sub>a</sub> (base) | 7.9<sup>a</sup> | 9.12<sup>b</sup> |
| f<sub>u,p</sub> | 0.64<sup>c</sup> | 0.89<sup>d</sup> |
| B/P | 1.08<sup>e</sup> | 0.55<sup>e</sup> |
| Permeability (10<sup>-6</sup> cm/s) | 26.3<sup>f</sup> | 23.8<sup>f</sup> |
| **Distribution<sup>g</sup>** |  |  |
| K<sub>p,adipose</sub> | 0.50 | 0.3 |
| K<sub>p,bone</sub> | 2.07 | 0.3 |
| K<sub>p,brain</sub> | 1.97 | 0.3 |
| K<sub>p,gastrointestinal tract</sub> | 5.49 | 0.3 |
| K<sub>p,heart</sub> | 6.79 | 0.3 |
| K<sub>p,kidney</sub> | 5.54 | 0.3 |
| K<sub>p,liver</sub> | 11.19 | 0.3 |
| K<sub>p,lung</sub> | 1.77 | 0.3 |
| K<sub>p,muscle</sub> | 2.50 | 0.3 |
| K<sub>p,pancreas</sub> | 4.19 | 0.3 |
| K<sub>p,skin</sub> | 0.50 | 0.3 |
| K<sub>p,spleen</sub> | 6.27 | 0.3 |
| **Metabolism** |  |  |
| CL<sub>total,iv</sub> (L/hr) | 83.5<sup>h</sup> | - |
| CL<sub>h</sub> (L/hr) | 75.3<sup>i</sup> | 0 |
| CL<sub>intrinsic</sub> (L/hr) | 520<sup>j</sup> | 0 |
| f<sub>m,M6G</sub> | 0.1<sup>k</sup> | - |
| **Excretion** |  |  |
| CL<sub>r</sub> (L/hr) | 8.24<sup>l</sup> | 10.84<sup>l</sup> |
| CL<sub>secretion</sub> (L/hr) | 39.3 (13.1×3)<sup>m</sup> | 43.5 (14.5×3)<sup>m</sup> |

f<sub>u,p</sub>, fraction unbound in plasma; B/P, blood-to-plasma ratio; K<sub>p</sub>, tissue-to-plasma partition coefficient for specific organ/tissue; CL<sub>total,iv</sub>, total body clearance after intravenous administration; CL<sub>h</sub>, hepatic clearance; CL<sub>intrinsic</sub>, metabolic intrinsic clearance; f<sub>m,M6G</sub>, fraction of
morphine metabolism to M6G; $CL_r$ , kidney clearance; $CL_{secretion}$ , unbound intrinsic kidney active secretion clearance at proximal tubule (clearance value of each proximal subsegment S1, S2, and S3)
<sup>a</sup>(21), <sup>b</sup>collected from www.drugbank.ca, <sup>c</sup>(16), <sup>d</sup>(17), <sup>e</sup>(18), <sup>f</sup>experimentally determined using VPT-MPS as described in the Methods, <sup>g</sup>predicted (22) and optimized as described in the Methods, <sup>h</sup>(25), <sup>ij</sup> calculated as described in the Methods, <sup>k</sup>(27), <sup>l</sup>predicted using mechanistic kidney model(10) as described in the Methods, <sup>m</sup>experimentally determined using VPT-MPS as described in the Methods

### Simulation of plasma morphine and M6G concentration-time profiles in healthy subjects and end stage kidney disease patients

All simulations were performed using MATLAB and Simulink platform (R2018a; MathWorks, Natick, MA) with intravenous administration and the same dose as reported in the corresponding clinical studies used as test set. For healthy subjects, to verify the morphine model, plasma morphine concentration-time profiles were simulated after intravenous administration of morphine and compared to the observed data from 2 test sets (28, 29) that were not used in model development. For M6G model verification, the plasma M6G concentration-time profiles were simulated after intravenous administration of M6G and compared to the observed data (24). Further, to verify the established parent-metabolite link between morphine and M6G, the plasma M6G concentration-time profiles were simulated as a metabolite after intravenous administration of morphine and compared to the observed data from 2 test sets (28, 29).

After verification of morphine and M6G models in healthy subjects, morphine and M6G disposition was also simulated in chronic kidney disease (CKD) patients using our published adaptive kidney model of progressive CKD (15). First, the kidney clearance of morphine and M6G was simulated across varying stages of CKD with GFR values ranging from 3 to 120 mL/min. The transporter-mediated active secretion of morphine and M6G was assumed to decrease proportionally with GFR in the simulations and the decrease in CL_r_ as a function of GFR was simulated. To simulate morphine and M6G systemic disposition in end stage kidney disease (ESKD) patients, the simulated kidney clearance of morphine and M6G at GFR = 3 mL/min was used, while all other parameters remain unchanged. The plasma morphine concentration-time profile and plasma M6G concentration-time profile as a metabolite of morphine were simulated after intravenous administration of morphine and compared to the observed data in ESKD patients (reported patient GFRs ranged from 0 to 5 mL/min) (13). All simulated plasma morphine and M6G concentrations were sampled from peripheral arm vein sampling site in the model (20) to match with the sampling site used in the clinical studies (test sets). To assess model performance, absolute average fold error was calculated using simulation results and observed data according to equation 10. The calculated absolute average fold error had to be within 2-fold (model acceptance criterion) for the simulation to be considered successful.

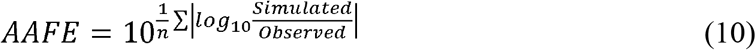

### Statistical Analysis

Statistical analysis was conducted in GraphPad Prism 7. A 2-tailed Student’s t test was employed to assess difference in concentrations of morphine or M6G in the PTECs effluents with and without treatment of inhibitor. *P* value less than 0.05 was considered significant.

## Results

### Construction of VPT-MPS and expression of transporters in VPT-MPS

Construction of the VPT-MPS for assessing tubular secretion of morphine and M6G is shown in Figure 1. The complete dual channel device is composed of an endothelial vessel, interstitial matrix, and an epithelial tubule, allowing the evaluation of the sequential processes of drug entry into and extravasation from the endothelial vessel, diffusion across the interstitial matrix and coordinated uptake and efflux of solutes across the tubular epithelium. Endothelial and epithelial tubular cells, human umbilical vein endothelial cells (HUVECs) and human primary proximal tubular epithelial cells (PTECs) were seeded in side-by-side channels of MPS devices, respectively, cultured under flow, and used for transport experiments. Immunocytochemical staining of PTECs in the VPT-MPS exhibited expression of organic anion transporter (OAT) 1 and 3 and organic cation transporter (OCT) 2 localized to basolateral membrane (Figure 1C).

### Transport of morphine and M6G across the epithelial tubule in the VPT-MPS

To evaluate secretory transport in the VPT-MPS, morphine and M6G were infused into the vascular channel in the presence and absence of an OAT and OCT inhibitor cocktail (1mM probenecid and 1 mM tetraethylammonium). Net clearances were calculated under steady state conditions (after 6-hours of infusion). Transport experiments were carried out in VPT-MPS from three different donors using devices with transepithelial electrical resistance (TEER) values of □150Ω*cm^2^ (30-32) and that demonstrated even, non-perturbed media flow characteristics. Data from a representative donor are shown in Figure 2. Calculated intrinsic transport and passive diffusion clearances for all the three donors tested in the study are shown in Table 1. Interestingly, efflux of morphine and M6G were markedly attenuated by transporter inhibitors with a 74.3% decrease in morphine intrinsic clearance (average of three donors, individual values varied from 54.1 to 86.1%) and of M6G by 63.6% (average of three donors, individual values varied from 17.1 to 87.1%), respectively. This suggests that active secretion, likely via OATs and OCT2, contributes to morphine and M6G kidney clearance.

**Figure 2.**
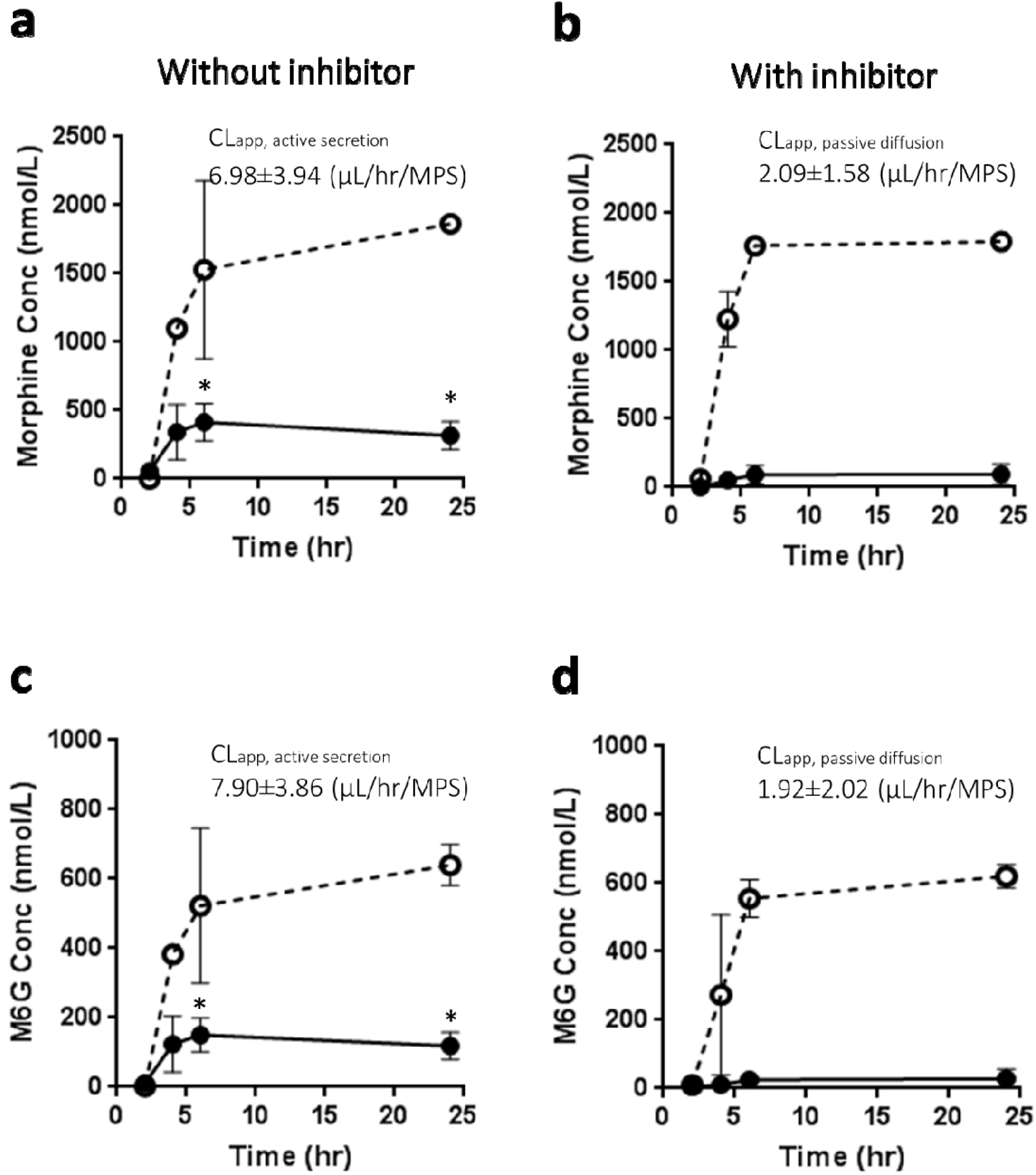
Transport of morphine and M6G by VPT-MPS. Concentration-time profiles of Morphine (a and b) and M6G (c and d) sampled from the respective effluents of endothelial vessel and epithelial tubule. Morphine and M6G were administered to vascular channel of VPT-MPS in the presence (b and d) and absence (a and c) of inhibitors and effluents from proximal tubule channel were collected. Dotted and solid line represents input in vascular concentration and output from epithelial tubule, respectively. Inset shows kinetic parameters of CL_app, passive diffusion_ and CL_app,active secretion_ calculated from MPS data in the presence and absence of inhibitors, respectively. Data represent mean ± SD from one representative donor. * indicates significant difference between the presence and absence of inhibitor (*p* < 0.05).

### Prediction of kidney clearance of morphine and M6G using VPT-MPS data and mechanistic kidney model

The kidney clearances of morphine and M6G were predicted using *in vitro* data from VPT-MPS and the previously published physiologically-based mechanistic kidney model (10). The experimentally determined CL_int secretion_, permeability, and predicted kidney clearances in three different donors are summarized in Table 1. The mean predicted kidney clearances of morphine and M6G using mechanistic kidney model and VPT-MPS data from three donors were 7.58 ± 2.53 L/h (range 4.8 to 9.7) and 9.45 ± 2.21 L/h (range 7.2 to 11.6), respectively. In comparison, the observed kidney clearances of morphine and M6G in human are 6.8-9.6 L/hr and 9.20-14.3 L/hr, respectively (28, 33, 34), suggesting that VPT-MPS together with the mechanistic kidney model generated successful predictions of kidney clearances of morphine and M6G with predicted values within 2-fold of the observed. Kidney clearances of morphine and M6G were also predicted using data from PTEC monolayers in 2D Transwell™, but the data from 2D cultures dramatically underpredicted the kidney clearance with mean predicted values of 2.46 ± 0.54 L/h and 4.11 ± 0.49 L/h for morphine and M6G, respectively (supplemental Table S4). Together, these results suggest that 3D VPT-MPS provides unique advantages compared to a 2D system in predicting kidney clearance of drugs and metabolites that have substantial active tubular secretion.

### Prediction of kidney clearance and systemic disposition of morphine and M6G in healthy subjects and end stage kidney disease patients using parent-metabolite full body PBPK model

The morphine (Figure 4 panel a and b) and M6G (Figure 4 panel c and d) drug models were independently verified using observed plasma morphine and M6G concentration-time data in healthy subjects after intravenously administering morphine (28, 29) or M6G (24). Then, the morphine-M6G parent-metabolite link was verified using observed plasma M6G concentration-time data after intravenous administration of morphine to healthy subjects (28, 29). The absolute average fold error values for morphine and M6G simulations (Figure 4) ranged from 1.18 to 1.33, all within the pre-defined 2-fold model acceptance criterion, suggesting successful model verification and high confidence on the model parameters in healthy subjects for both morphine and M6G.

**Figure 3.**
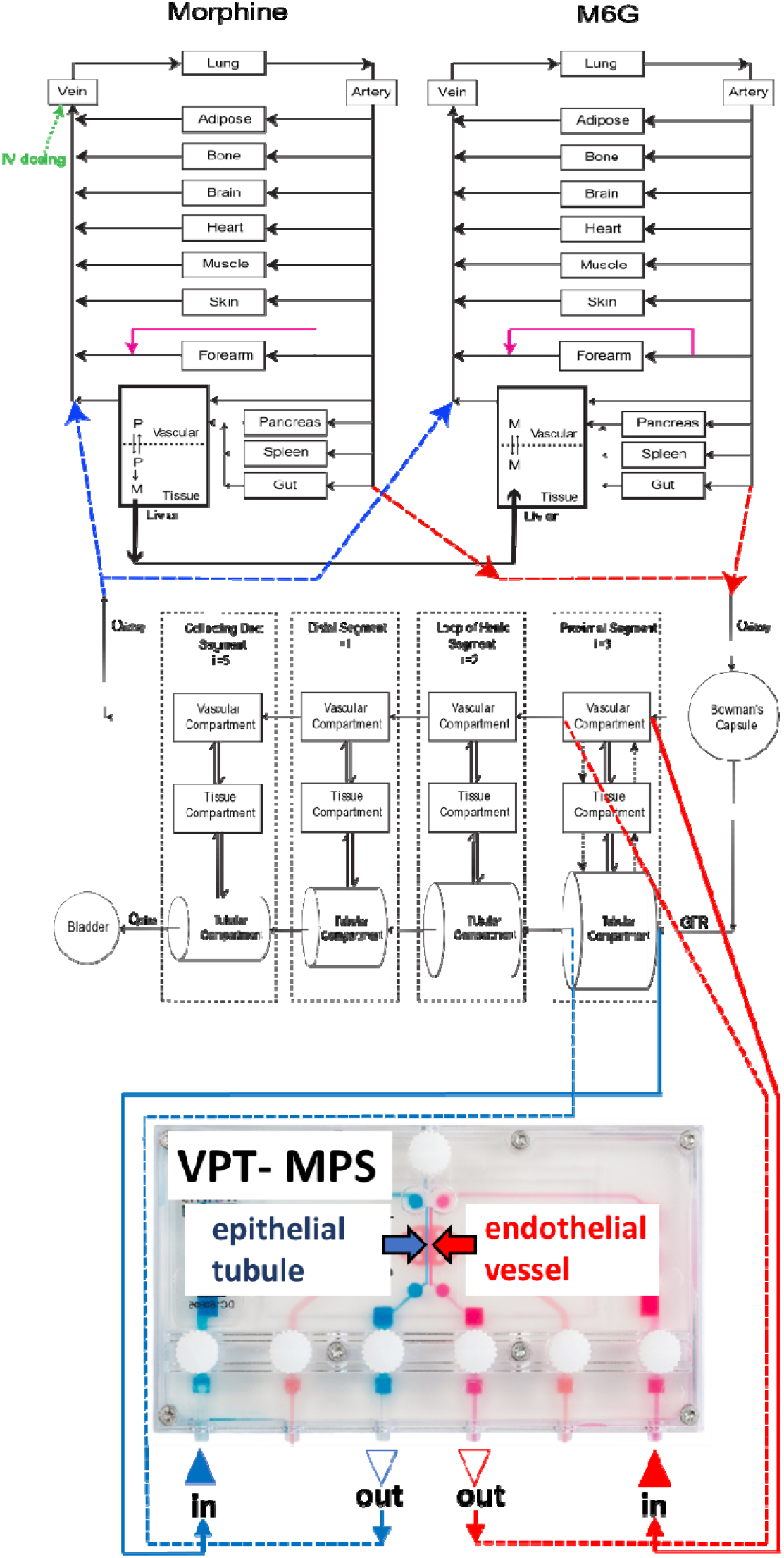
Structure of the mechanistic morphine-M6G parent-metabolite full body PBPK model. Schematic presentation of the physiologically based parent-metabolite pharmacokinetic model that incorporates a mechanistic kidney model, adapted from previous publication (11). The transporter-mediated active secretion or active reabsorption is shown in black dotted arrows. The bidirectional pH-dependent passive diffusion is shown in double arrows. Lower panel shows VPT-MPS used for evaluation of active transport and passive diffusion of Morphine and M6G, where endothelial vessel and epithelial tubule were shown in red and blue, respectively. Q_kidney_, kidney blood flow; Q_urine_, urine formation flow; GFR, glomerular filtration rate; i, the number of subsegments.

**Figure 4.**
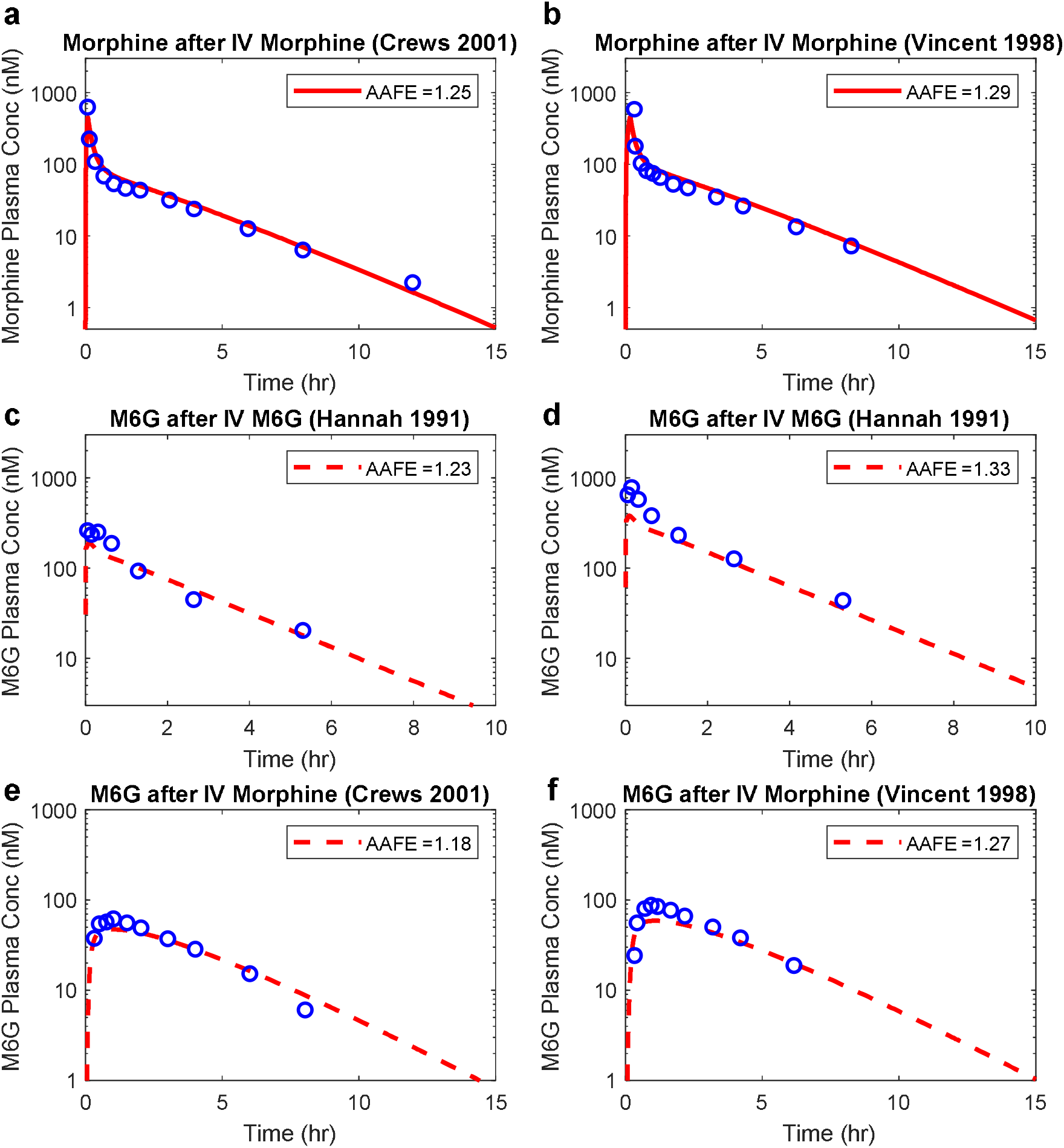
Simulation of morphine and M6G plasma concentration-time profiles in healthy subjects. Plasma morphine concentration-time profile was simulated after intravenous dosing of morphine (shown in red solid curves) in comparison to the observed data in healthy subjects from two independent studies (28, 29) (panel a and b). Plasma M6G concentration-time profile was simulated after intravenous dosing of M6G (shown in red dashed curves) in comparison to the observed data in healthy subjects from two different doses: 30 µg/kg (panel c) and 60 µg/kg (24) (panel d). Plasma M6G concentration-time profile was simulated as a metabolite (shown in red dashed curves) after intravenous dosing of morphine in comparison to the observed data in healthy subjects from two independent studies (panel e and f) (28, 29). All simulation results are shown in red and all observed data are shown in blue open circles. The calculated absolute average fold error value for each panel is shown in the inset.

After model verification, the model was extrapolated to investigate disposition of morphine and M6G in CKD patients. First, the kidney clearances of morphine and M6G were simulated across multiple stages of CKD ranging from glomerular filtration rate (GFR) estimate of 3 mL/min to 120 mL/min (Figure 5 panel a). The simulations showed that M6G kidney clearances are 10-30% higher than morphine kidney clearances throughout CKD progression (Figure 5 panel a). Then, the CKD model of morphine and M6G was verified using observed plasma morphine and M6G concentration-time profiles after intravenous administration of morphine in ESKD patients (13). The absolute average fold error values for CKD simulations (Figure 5 panel b and c) ranged from 1.22 to 1.30, all within the pre-defined 2-fold model acceptance criterion, demonstrated successful model extrapolation of morphine and M6G from healthy subjects to CKD patients. This indicates that the *in vitro* results obtained from VPT-MPS can be translated to *in vivo*, through an appropriate *in silico* framework.

**Figure 5.**
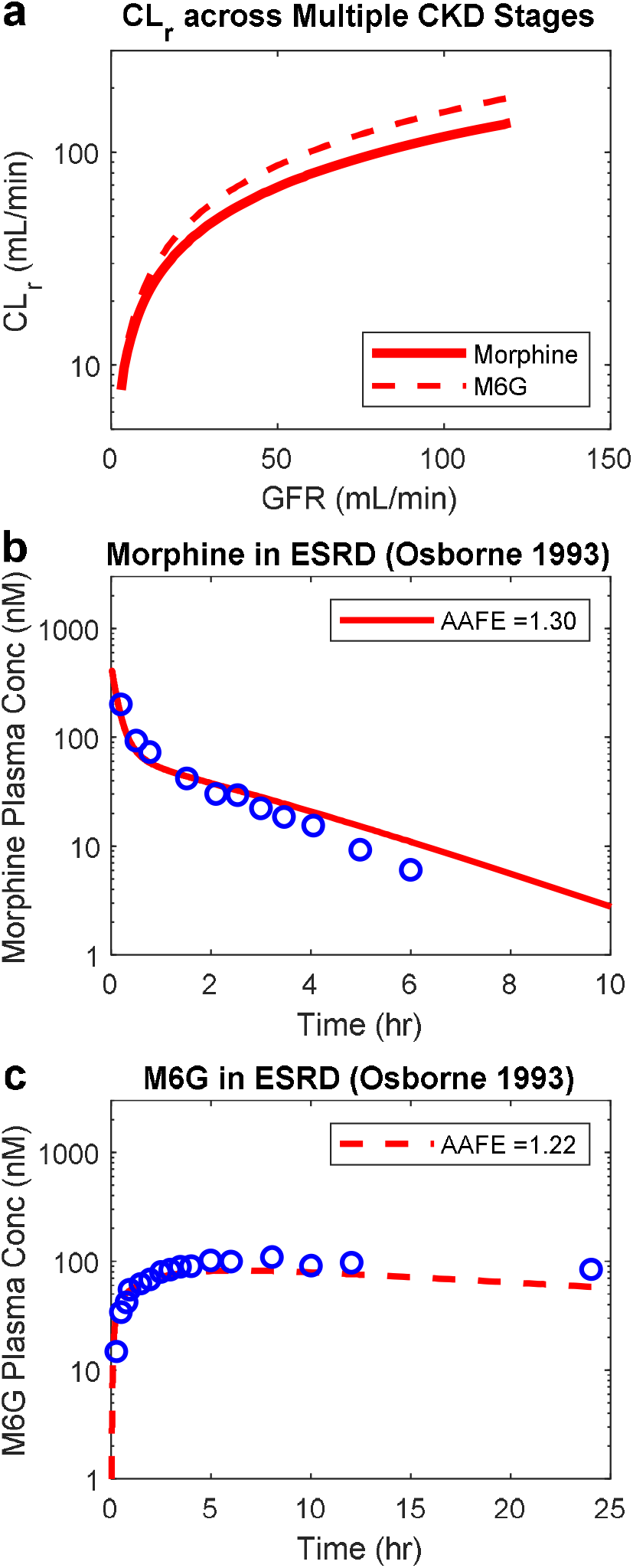
Simulation of morphine and M6G kidney clearance and plasma concentration-time profiles in chronic kidney disease patients. The kidney clearance (CL_r_ in mL/min) of morphine (red solid curve) and M6G (red dashed curve) was simulated at multiple stages of chronic kidney disease (CKD) reflected by varying glomerular filtration rate (GFR in mL/min) using novel *in silico* adaptive CKD model (15) together with experimentally determined permeability and active secretion via VPT-MPS (panel a). The plasma morphine concentration-time profile was simulated after intravenous dosing of morphine (shown in red solid curves) in end stage kidney disease (ESKD) patients in comparison to the observed data (13) (panel c). The plasma M6G concentration-time profile was simulated as a metabolite (shown in red dashed curves) in end stage kidney disease (ESKD) patients after intravenous dosing of morphine in comparison to the observed data (13) (panel c). All simulation results are shown in red and all observed data are shown in blue open circles. The calculated absolute average fold error value for each panel is shown in the inset.

**Figure 6.**
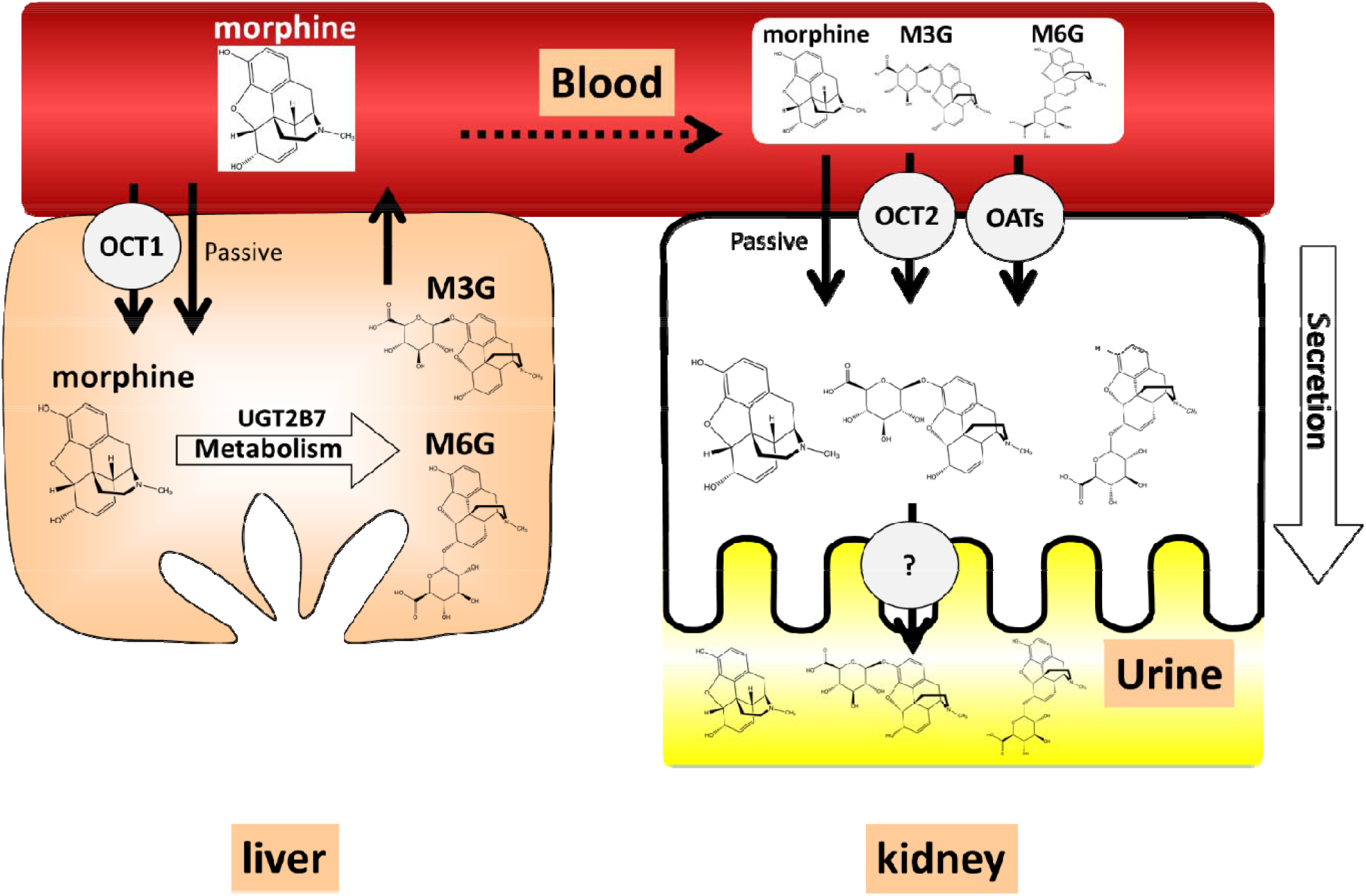
Structure of morphine and metabolites, schematic diagram of hepatic/kidney disposition of morphine and metabolites. Morphine is taken up into the liver via OCT1 and metabolized by UGT2B7 into glucuronide-con ugated metabolite, M3G and M6G. Morphine, M3G and M6G are subsequently transported out of liver into blood and taken up into the kidney by passive diffusion and active uptake by OATs (OAT1/OAT3) and OCT2 and undergo kidney secretion.

## Discussion

With the increased risk of opioid overdose and adverse events in people with CKD (12), there is a growing need for accurate prediction of opioid disposition in sensitive populations to mitigate the risk of opioid overdose, dependence and addiction. In this study, we hypothesized that the VPT-MPS, in concert with a mechanistic modeling approach, can predict the disposition of morphine and its active metabolite M6G in healthy subjects and patients with kidney impairment. Passive permeability and active secretion data were generated using the VPT-MPS system. The *in vitro* data were incorporated into a mechanistic kidney model-integrated parent-metabolite full body PBPK model to predict kidney clearance and systemic disposition of morphine and M6G in both healthy subjects and CKD patients. Our results show that this translational approach can successfully predict CL_r_ and plasma concentration-time profile with absolute average fold error < 1.5 in both healthy subjects and CKD patients, suggesting that the VPT-MPS together with PBPK modeling has the potential to bridge the *in silico* and *in vivo* gap.

Both morphine and M6G are pharmacologically active (33-35); M6G in particular and morphine to some degree are actively secreted into urine with kidney clearance exceeding plasma unbound fraction multiplied by glomerular filtration rate (33). Therefore, quantitative evaluation of the contribution of active secretion to the kidney clearance of both morphine and M6G is essential to predict the impact of attenuated kidney function on kidney clearance and subsequent systemic disposition in CKD patients. Yet, no relevant information is currently available on specific secretory clearance in the kidney, although the role of organic cation transporters (OCTs) in the hepatic uptake is recognized (36, 37). The present study utilized a VPT-MPS to evaluate the kidney secretion of morphine and M6G and to predict kidney clearance of these compounds in humans.

While a MPS can better predict CL_r_ of drugs in humans by recapitulation of *in vivo* physiology, *in vitro* systems alone cannot directly determine the intrinsic clearance of tested drugs. To predict CL_r_ and systemic disposition, additional methodologies are necessary to scale intrinsic clearances (generated with *in vitro* experiments) to *in vivo* organ level clearances. To this end, IVIVE has been widely accepted approach, where *in vitro* data are directly extrapolated to *in vivo* parameters by incorporating a scaling factor (2) based on analysis of several index compounds. However, this approach does not consider complex human physiology, resulting in inaccurate prediction. To bridge the *in vitro* and *in vivo* gap, mathematical models have been created that extrapolate *in vitro* data to *in vivo* PK parameters by incorporating drug-related parameters (e.g. passive permeation and active transport) together with system-specific parameters (e.g. tubular flow rates and pH values) (38, 39). In the present study, we employed a VPT-MPS platform coupled with mathematical modeling to predict human kidney clearance and plasma pharmacokinetics.

In the VPT-MPS experiments, following infusion of morphine or M6G via the vascular channel, efflux of morphine and M6G into the proximal tubule channel effluent was observed in a time-dependent manner. Moreover, this efflux was inhibited by coadministration of tetraethylammonium (OCT inhibitor) and probenecid (organic anion transporter (OAT) inhibitor) (Figure 2), suggesting that functionality and expression of transporters are maintained in the VPT-MPS. This conclusion was supported by immunohistochemical localization of transporters (Figure 1). Moreover, we contend that the contribution of transporter(s) involved in the active kidney secretion of morphine and M6G can be quantitatively evaluated in our VPT-MPS.

The contribution of OATs to kidney secretion of M6G has been demonstrated in clinical studies (40), in agreement with our observations. While active secretion of morphine and M6G was confirmed in VPT-MPS, it should be noted that we observed significant variability in the active secretion of morphine and M6G between PTECs donors with CL_int, active secretion_ values ranging from 1.33 to 20.9 µL/hr/VPT-MPS for morphine and 1.29 to 22.9 µL/hr/VPT-MPS for M6G, respectively (Table 1). Although the origin of this variability is unclear, possible causes include inter-donor differences of expression of transporters (as reported by Nozaki et al who demonstrated marked variabilities in expression and activity of OATs in kidney slices (1)), and warm ischemic conditions during the nephrectomy process (which may impact expression and function of transporters). Indeed, Schneider et al. have demonstrated that rat Oat1 and Oat3 expression and function were significantly impaired by acute ischemia reperfusion injury rat models (41).

To extrapolate *in vitro* data to *in vivo* pharmacokinetics, we incorporated experimental results obtained from VPT-MPS into a state-of-art physiologically-based mechanistic kidney model (10). As shown in Table 1, the mean kidney clearances predicted from VPT-MPS data together with mechanistic kidney PBPK model were 7.58 ± 2.53 L/h for morphine and 9.45 ± 2.21 L/h for M6G, respectively, which both fall within 1.5-fold of the observed mean kidney clearances (28, 33, 34). In contrast, 2D Transwell™ dramatically underpredicted kidney clearance due to a lack of active secretion in 2D culture (Table S4), likely due to minimal transporter expression as confirmed by negligible active transport of *para*-amminohippuric acid, a probe substrate of OAT1 and OAT3 (Table S3). Taken together, VPT-MPS coupled with a mechanistic kidney PBPK model that incorporates glomerular filtration, active secretion and passive reabsorption successfully predicted human kidney clearance of morphine and M6G.

In addition to kidney clearance, systemic disposition of morphine and M6G in healthy subjects and CKD patients was also simulated, as patients with kidney impairment may experience greater exposure and prolonged half-life of morphine and M6G, leading to increased risk of opioid overdose (13). Using a parent-metabolite full body PBPK model coupled with the VPT-MPS data-populated mechanistic kidney model, the plasma concentration–time profiles of morphine and M6G in healthy subjects were successfully simulated with all absolute average fold error values less than 1.5. Further, the model was successfully extrapolated to simulate morphine and M6G systemic disposition in CKD patients by modification of GFR, tubular flow rate/passive reabsorption, and transporter-mediated active secretion, with all absolute average fold error values less than 1.5, demonstrating successful model extrapolation from healthy subjects to CKD patients.

It should be noted that in the present study, we calculated the hepatic clearance of morphine to simulate systemic disposition of morphine by subtracting the experimentally determined morphine kidney clearance from the observed morphine systemic clearance after intravenous administration (25). However, in an assessment of impact of hepatic metabolism on systemic exposure of morphine and M6G in addition to kidney elimination, use of systems such as liver-kidney linked microphysiological systems are likely to be useful, enabling replication of sequential hepatic metabolism/elimination (in our case, conversion of morphine to M6G) and kidney elimination. This approach will require an *in vitro* MPS that recapitulates *in vivo* human physiology that considers the physiological differences between the two organs (e.g., differential rates of blood flow). This is especially important when two or more organs linked in MPS are used for quantitative assessment of pharmacokinetic, pharmacodynamic or toxicodynamic studies (42).

In this study, parameters governing kidney transport of morphine and M6G across proximal tubules were obtained from VPT-MPS experiments, which were then used to inform a mathematical PBPK model that simulated systemic exposure of morphine and M6G in subjects with normal and impaired kidney function. Given that CKD patients experience altered pharmacokinetics (PK), accurate prediction of PK is critical for safe and effective use of medications in this population. To address these challenges, several attempts have been made to simulate PK in CKD patients using PBPK modeling (43, 44). However, there are limitations in previously published studies, particularly in the prediction of CL_r_. For example, active secretion was attributed solely to organic anion transporters (OATs) and all compounds tested were organic anions; thus generalizability is limited for compounds that are substrates of multiple transporters (44). In addition, previous studies tested only compounds where kidney clearance significantly contributed to total clearance; it is unknown whether this modeling approach can be applied to compounds with a lower contribution of active kidney tubular secretion to overall kidney clearance, or drugs which undergo elimination by both kidney and hepatic pathways (43, 44). In contrast, our current approach utilizes data generated from MPS, which enables estimation of active kidney secretion regardless of the mechanism of transport of compounds of interest. Also, applicability of our model to simulate human PK has also been validated for compounds for which active kidney secretion is limited and kidney elimination is sensitive to urine pH and flow (10, 11) in addition to compounds with significant active transport. In this study, we assumed that the transporter-mediated active secretion of morphine and M6G decreases proportionally with GFR in accordance with the intact nephron hypothesis (45). However, several groups have reported that alterations in kidney transport protein expression/activity and inhibition of kidney disposition by circulating uremic solutes occur in CKD patients (46, 47); these possibilities should be taken into consideration by conducting *in vivo* experiments to determine appropriate scaling combined with *in vitro* experiments to investigate inhibitory effect of various uremic solutes on transporters activity and variable expression of transporters in patients across various stages of CKD.

In conclusion, we have presented a novel translational strategy that couples a VPT-MPS with mechanistic PBPK modeling for quantitative prediction of the kidney disposition of drugs. The VPT-MPS recapitulates critical aspects of kidney physiological function *in vivo*, and allows evaluation of active secretory clearance of morphine and M6G with sustained maintenance of transporter protein expression. The mechanistic kidney model and the parent-metabolite PBPK model successfully incorporated the VPT-MPS data and accurately predicted kidney clearance and plasma concentration-time profiles of morphine and M6G in both healthy subjects and CKD patients, indicating that the *in vitro* results obtained from VPT-MPS can be translated to *in vivo* parameters, via an appropriate *in silico* framework. Taken together, we have shown that the bottom-up approach can predict opioid disposition in CKD patients and potentially mitigate the risk of overdose in those sensitive populations. We contend that this approach can be further extrapolated to a variety of drugs and investigational compounds to *a priori* understand kidney and systemic disposition in vulnerable patient populations.

## Author contributions

T.I., W.H., and S.S. designed, conducted and analyzed *in vitro* MPS experiments and modeling and simulation works. D.W.H conducted imaging studies of MPS experiments. C.K.Y, N.I., E.J.K. and J.H. supervised and managed the experiments. All the authors contributed to writing the manuscript.

## Acknowledgments

None

## Supplementary Materials Table of Contents

### Materials and Methods

Supplemental Figure 1. Metabolism and transport of morphine using liver MPS

Supplemental Table 1. Donor information on kidney tissues used for isolation of proximal tubule epithelial cells.

Supplemental Table 2. Transport of morphine and M6G by VPT-MPS

Supplemental Table 3. TEER and Papp values for the bidirectional transport of Morphine, M6G and [^3^H]PAH across PTECs and MDCK monolayers after 1.5 hour incubation in conventional 2D Transwell™.

Supplemental Table 4. Prediction of kidney clearance of morphine and M6G from PTECs permeability study data

